# Protease-activated receptor 2 links protease activity with calcium waves during egg activation and blastomere cleavage

**DOI:** 10.1101/2024.05.14.594074

**Authors:** Jiajia Ma, Tom J. Carney

**Affiliations:** Lee Kong Chian School of Medicine, Experimental Medicine Building, Yunnan Garden Campus, 59 Nanyang Drive, Nanyang Technological University, 636921, Singapore; Institute of Molecular and Cell Biology (IMCB), A*STAR (Agency for Science, Technology and Research), 61 Biopolis Drive, Proteos, 138673, Singapore

**Keywords:** Protease-activated receptor 2, Proteinase-activated receptor 2, Par2, zebrafish, egg activation, oocyte, calcium, serine protease, fertilisation, calcium signalling

## Abstract

Successful initiation of animal development requires activation of the egg immediately prior to fusion of gamete pronucleii. In all taxa, this is initiated by waves of calcium transients which transverse across the egg. Calcium waves also occur at cleavage furrows during later blastula cytokinesis. Calcium is released from the endoplasmic reticulum through activation of 1,4,5-trisphosphate (IP_3_) receptors. Only a subset of the mechanisms employed to generate IP_3_ during vertebrate egg activation are defined, with strong evidence that other critical mechanisms exist. Serine proteases have been long implicated in egg activation and fertilisation. Here we report that treatment of zebrafish eggs with serine protease inhibitors leads to defective calcium wave propagation and failed egg activation. We further show that mutation of zebrafish Protease-activated receptor 2a (Par2a) also results in severe disruption of egg activation, leading to failed chorion elevation and ooplasmic segregation. Milder *par2a* mutants progress further, but then show abnormal blastomere cleavage. We observed that *par2a* mutants show decreased amplitude and duration of calcium transients. Restoring Ca^++^ or direct injection of IP_3_ ligand rescues egg activation aborted by either serine protease inhibitor treatment or by mutation of Par2a. We thus show that serine protease activity is a critical regulator of IP_3_ and subsequent calcium wave amplification during zebrafish egg activation, and link this to intracellular calcium release via the protease receptor, Par2a. This constitutes a novel signalling pathway critical for successful fertilisation.

**Significance Statement:** Both sperm and egg must undergo a series of important steps to become competent for successful fertilisation. Defining these steps is central to our understanding of reproductive biology and our ability to improve fertility treatments. As the process of gamete maturation and fertilisation has highly conserved principles across the animal kingdom, there are also important implications for aqua- and agriculture. One of the first signalling events of your life leads to the release of bursts of calcium in the egg. We know the importance of this for fertilisation but have only a partial picture of how this occurs. Our work here, using fish genetics, identifies a new signalling pathway regulating these first important flashes of calcium in the egg.

## Introduction

Fertilisation culminates in the fusion of sperm and egg pronuclei to generate a zygote. Prior to this, both gametes must undergo a number of maturation and activation steps to achieve fertilisation competence (1). The final activation process of the oocyte is termed egg activation, which occurs upon, or immediately prior to, fertilisation. Egg activation has broad conservation across animals and achieves largely comparable outcomes in all species (2). These include exocytosis of cortical granules, the cortical reaction, formation of the pronucleus, resumption of meiosis and extrusion of the second polar body. These are essential for blocking polyspermy and for subsequent development to proceed. In fish species, the release of cortical granules during activation elevates the chorion and creates a perivitelline fluid filled space between the chorion and the egg. There is also a significant re-organisation of the egg cytoplasm which is initially intermingled with lipid rich yolk granules in arrested fish eggs. Upon activation, actin-myosin contractions squeeze this ooplasm away from the yolk to the animal pole where it forms the blastodisc (3, 4).

Whilst in mammals egg activation is initiated by binding of sperm to the egg, in fish, egg activation occurs upon contact of the eggs with the water during spawning, and thus fish egg activation does not require fertilisation and can occur parthenogenetically (5). Spontaneous activation of zebrafish eggs following spawning into water can be blocked, however, by incubation in fish ovarian fluid (6). The critical factor imparting this property has been proposed to be a serpin-type protease inhibitor which is found at high levels in zebrafish ovarian fluid (7). Furthermore, treatment of loach and carp eggs with protease inhibitors also prevented spontaneous activation (8), although it is unclear how protease inhibitors block egg activation.

Intracellular calcium has a fundamental and conserved role in driving egg activation, fertilisation, and early embryo development across both vertebrates and invertebrates (2, 5, 9-11). The dynamics of calcium varies in the eggs of different species. In most invertebrates, fish, and amphibia, egg activation is driven by a single Ca^++^ wave that transverses the egg, whilst in mammals, fertilisation initiates multiple oscillatory waves (5, 12). Irrespective of initiation mode or Ca^++^ wave pattern, assays in multiple species including sea urchins, *Xenopus*, fish and mammals, have demonstrated the second messenger, inositol-1,4,5-trisphosphate (IP_3_) releases intracellular Ca^++^ in eggs via binding to IP_3_ receptors on the endoplasmic reticulum. These IP_3_ generated Ca^++^ waves are sufficient to activate eggs (2, 13-16).

In amphibia and fish, distinct Ca^++^ transients are later observed extending laterally along the nascent division plane of cleavage furrows prior to blastomere cytokinesis (17-19). These transient waves of Ca^++^ and are associated with furrow positioning, propagation and deepening (20). As with egg activation, there is a requirement for IP_3_ in these transients although there is evidence that each Ca^++^ transient may have distinct regulation mechanics (21-23).

The varied mechanisms and environments initiating egg activation across the animal kingdom and the diverse patterns of Ca^++^ waves employed therein and subsequently during blastomere cleavage, indicates that there are highly likely to be multiple Ca^++^ regulating mechanisms. As IP_3_ is generated by cleavage of PIP_2_ through the activity of Phospholipase C (PLC) family members, it is likely that PLC regulation plays a central role in egg activation. The first Ca^++^ transient in mouse is initiated by release of phospholipase C zeta (PLCζ) from the sperm into the egg cytoplasm (24). In sea urchin and xenopus, FAK and Src-family protein tyrosine kinases contribute to Ca^++^ release during egg activation through activation of PLCγ, although Src family kinases are entirely dispensable for Ca^++^ release in the mouse and have only a partial effect in zebrafish (25-29).

In addition to kinases, PLCs are also activated by the heterotrimeric G protein alpha subunit, Gq, which in turn is activated by certain G protein-coupled receptors (GPCRs). Protease Activated Receptors are a family of GPCRs activated by extracellular protease activity, through unmasking of a tethered ligand which binds to the receptor intra-molecularly activating downstream signalling (30). For example, Par2 is activated by trypsin-like proteases and has been shown to activate Gq and in turn PLCβ, thus releasing calcium in a number of cell types and contexts (31-33).

Here we show that zebrafish egg activation is sensitive to serine protease inhibitors, and identify that maternal mutants of the serine protease responsive receptor, Par2a, have broad egg activation defects due to altered calcium waves. This identifies a novel essential regulator of calcium during the activation of vertebrate eggs.

## Results

### Serine protease inhibition stalls egg activation

We wanted to determine if zebrafish egg activation was sensitive to protease inhibitors as shown for the loach and carp (8). Eggs were squeezed from WT zebrafish females into Hank’s Solution containing sperm. This solution also contained 0.5% BSA, which is known to hold eggs in an inactivated state and block fertilisation (34). Eggs were then activated by diluting out of the BSA through addition of excess E2 medium, thus permitting fertilisation to proceed. Eggs showed immediate activation, rapidly raising chorions and displaying a prominent blastodisc after 30 minutes, with over 90% subsequently showing normal cell division indicating successful fertilisation (**Fig. 1A**). Addition of 5mg/ml of the peptide-based serine protease inhibitor, Aprotinin, to the E2 medium however, resulted in all eggs remaining fully inactivated even after 30mins, showing neither chorion elevation nor blastodisc formation and thus no cell division (100%, n>200 eggs; **Fig. 1B**). Presence of sperm made no difference to egg activation block by Aprotinin, indicating that Aprotinin is disrupting egg activation and not fertilisation.

**Figure 1:**
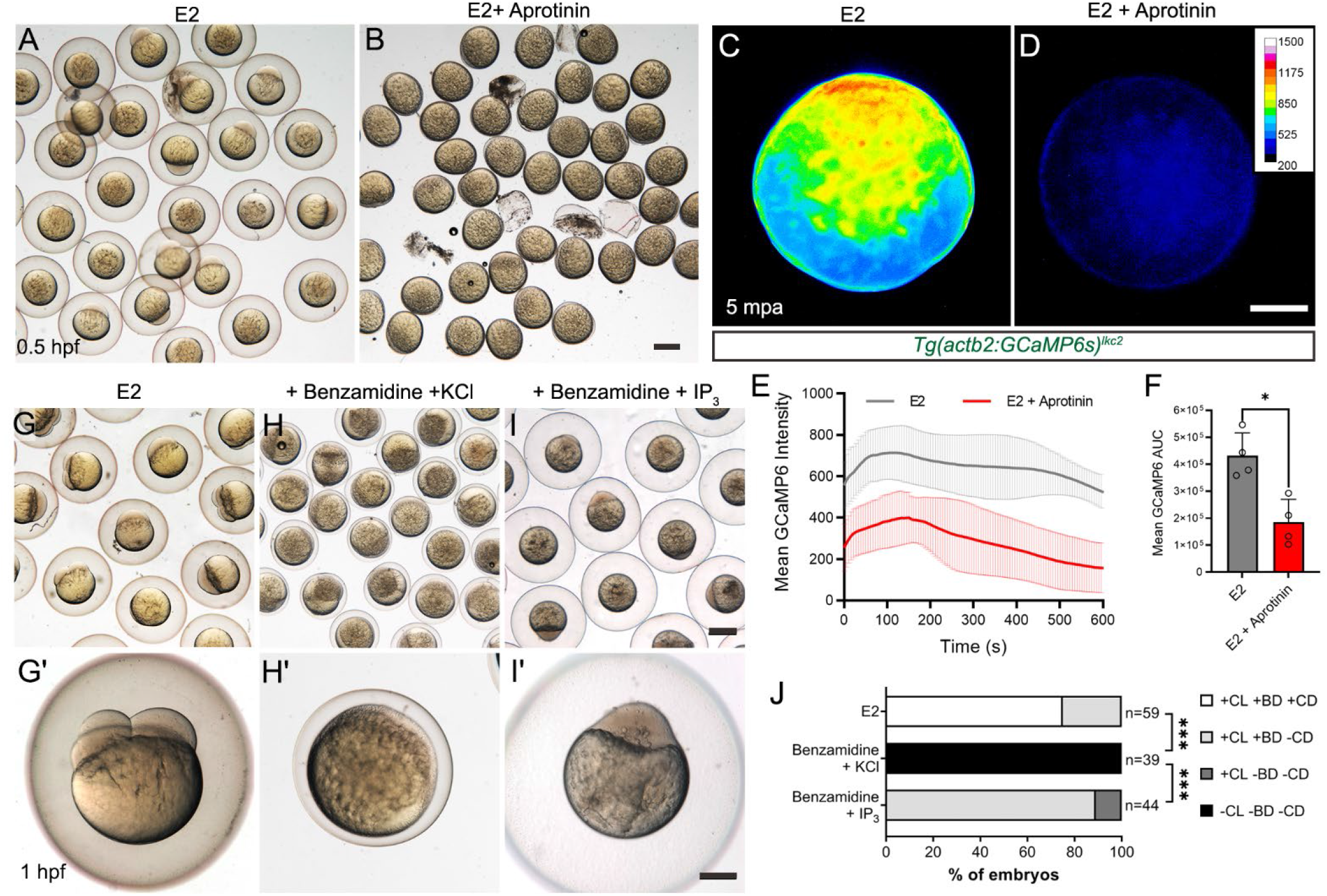
Serine protease inhibitors cause egg activation defects and reduced calcium waves. **A-B:** Eggs fertilised *in vitro* in E2 medium with (B) or without (A) 5mg/ml Aprotinin. **C-D:** Fluorescent images of eggs harvested from a *Tg(actb2:GCaMP6s)*^*lkc2*^ female, indicating Ca^++^ levels at 5 minutes post activation (mpa) in E2 (C) or in 5mg/ml Aprotinin (D). **E:** Changes in GCaMP6s intensity over 10 minutes of *Tg(actb2:GCaMP6s)*^*lkc2*^ eggs activated with E2 (black) or 5mg/ml Aprotinin (red). **F:** Corresponding Area Under Curve of GCaMP6 intensity levels presented in (E) following egg activation. n=4; Mann-Whitney test; * = p<0.05. **G-I’:** Low (G, H, I) and high (G’, H, I’) magnification images of embryos fertilised *in vitro* in E2 (G-G’), or in E2 with 40mg/ml Benzamidine HCl treatment and subsequently injected with 0.7mM KCl (H-H’), or in E2 with 40mg/ml Benzamidine HCl treatment and injected with 10mM IP_3_ in KCl (I-I’). **J:** Proportion of embryos showing egg activation phenotypes and blastomere division. Benzamidine incubation aborts egg activation, which can be reactivated by IP_3_ injection. Key: CL: Chorion Lift, BD: Blastodisc, CD: Cell Division +: Present, -: Absent. Chi-squared analyses; *** = p<0.001. Scale bars: B, I = 500µm; D, I’ = 200µm.

### Serine proteases activate eggs via IP_3_ mediated calcium release

To determine if protease inhibition altered calcium dynamics during egg activation, we employed the *Tg(actb2:GCaMP6s)*^*lkc2*^ transgenic line (32, 35). The *β-actin* promoter is active in oocytes and thus the GCaMP6s reporter protein is deposited in eggs. We activated eggs from this line in E2 with or without Aprotinin and visualised fluorescence. Ca^++^ transients in Aprotinin treated eggs were shorter and less pronounced compared to E2 treated controls (**Fig. 1C-D; Supplementary Movie 1**). Quantification over the first 10 minutes following Aprotinin treatment showed a significant reduction in amplitude and duration compared to E2 activated eggs (**Fig. 1E, F**). Similar results were obtained using Benzamidine HCl, a small molecule inhibitor of trypsin-like serine proteases. Benzamidine reduced egg activation during *in vitro* fertilisation, although less potently than Aprotinin, with eggs showing partial chorion lifting and small blastodisc formation (**Fig. 1G, G’, H, H’**; **Fig. S1A-B**). Benzamidine also significantly reduced duration of Ca^++^ transients during egg activation compared to controls (**Fig. S1C-E, Supplementary Movie 2**). Thus, the activity of serine proteases, most likely trypsin-like serine proteases, are necessary for generation of normal calcium waves and egg activation in zebrafish.

Injection of Inositol 1,4,5-trisphosphate (IP_3_) can activate eggs of many species (2). We asked if addition of IP_3_ could reactivate eggs arrested by serine protease inhibition. We injected 20pmol of IP_3_ into eggs treated with Benzamidine and observed robust restoration of egg activation, with successful chorion lifting and formation of a blastodisc compared to Benzamidine treated eggs injected with KCl injection solution only (**Fig 1G, G’, H, H’, I, I’, J**). However, these eggs were unable to undergo subsequent cell division, despite being derived from IVF, suggesting a serine protease is required for subsequent cell division in a process independent of IP_3_. Thus, serine protease activity is required for egg activation through a Ca^++^, and likely an IP_3_ dependant process.

### Maternal *par2a* mutants display defects in egg activation and blastomere cleavage

How IP_3_ is generated in eggs to initiate activation appears to be species specific. It is unclear what generates IP_3_ and how this may involve proteases. In zebrafish, Par2b has been implicated in IP_3_ formation and intracellular Ca^++^ release in the epidermis following activation by the serine protease Matriptase1a (32, 36). As *par2a* and *par2b* are also both maternally expressed (37), we hypothesised that they may link serine protease activity with IP_3_ and Ca^++^ generation during egg activation.

We generated *par2a* and *par2b* mutants through CRISPR/Cas9 targeting. CRISPR guide RNA was designed to target the 2^nd^ extracellular loop of the receptor. In all alleles, indels led to a frame shift leading to premature protein termination before the fifth transmembrane domain, and before the third intracellular and extracellular loops. The predicted protein in all alleles thus also lack the C-terminal tail (**Fig. S2A-C**). Zygotic homozygous mutants for all three *par2a* alleles (**Fig. S2A**) were adult viable with no overt phenotype. Female homozygous *par2a* mutant adults, however, gave rise to eggs that displayed a defect in activation, reminiscent of the activation phenotype seen in eggs treated with serine protease inhibitors. Thus, both activated eggs and embryos derived from natural spawning of *par2a*^*-/-*^ females with WT males showed significantly impaired chorion elevation, and defective blastodisc formation compared to WT crosses or activated WT eggs (**Fig. 2A; Table S1**). Expressivity was variable. Severe mutant embryos showed only minimal chorion elevation and no blastodisc formation, whilst milder mutant embryos displayed only moderate chorion lifting and partial blastodisc formation, however this blastodisc was often unstable and showed regions of partial collapse (**Fig. 2A Fig. S3D**). The effects are not due to simple delay, as even by 1.5 hours post activation, severely affected eggs remain with limited chorion elevation and no blastodisc (**Fig. S3A-F; Supplementary Movie 3**). In more mildly affected fertilised eggs, some blastodiscs do form of sufficient size that they initiate blastomere cleavages, however resulting daughter cells are of unequal sizes, ultimately leading to failure to form normal blastomere tiers. (**Fig. S4A-B; Supplementary Movie 3**). Blastomeres show reduced compaction and adherence to each other, and some detach into the perivitelline space (**Fig. S4A, B**). All detached cells appear to retain nuclei (**Fig. S4C**). Although mild *par2a* mutant embryos are able to generate blastomeres, less than 1% successfully complete gastrulation, and those that do have severely affected axes and do not survive beyond 5dpf (**Table S1**). All alleles were identical in ability to generate severe or mild phenotypes. We found that the generation of mild or severe phenotypes was a property of each mother, not of each allele. Thus, different females of the same allele will consistently generate either severe or mildly affected clutches and there was a consistent severity within clutches (**Fig. S4D**). To test if this variability was due to compensation by *par2b*, we made a *par2b* CRISPR allele which created an indel 303bp downstream of the CRISPR binding site and terminated the protein before the fifth transmembrane domain (**Fig. S2B, C**). This allele of *par2b* was homozygous viable, and eggs from mutant females developed normally. Double *par2a; par2b* mutant females still showed variable expressivity in egg activation phenotype, with no additive severity induced by combined loss of both paralogues (**Table S1**). This indicates that compensation by *par2b* does not account for variable expressivity, and that *par2a*, but not *par2b*, is crucial for egg to embryo transition.

**Figure 2:**
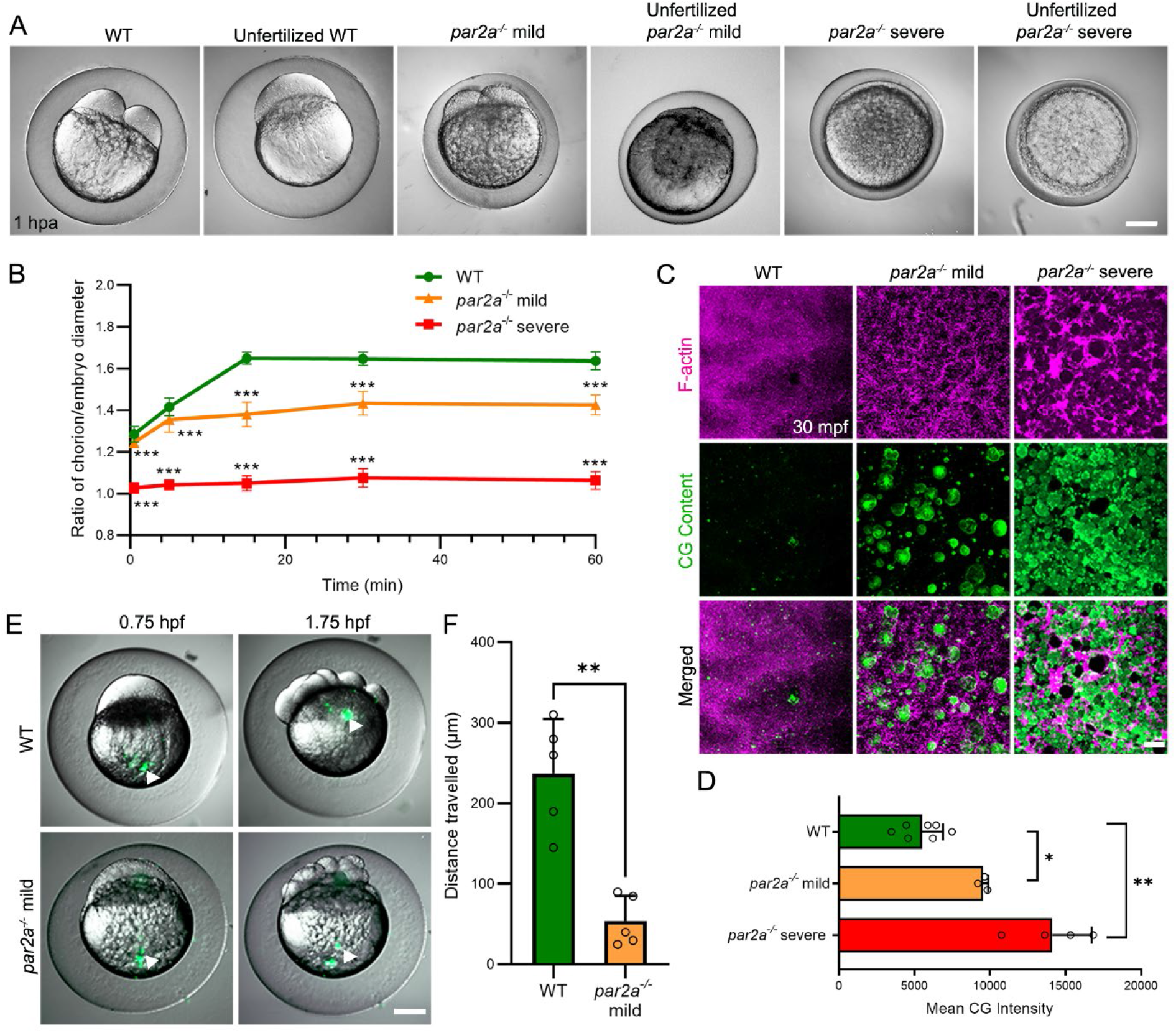
Maternal *par2a* mutant embryos display impaired egg activation events and cell division defects. **A:** Nomarski images of naturally fertilized and unfertilised, activated eggs from WT, mild and severe *par2a* mutant females. Chorion elevation and blastodisc formation are variably disrupted. **B:** Changes in ratio of chorion lift to embryo size of naturally fertilised WT, mild and severe *par2a* mutant eggs at different time-points over an hour post activation, n=51; t-test; *** = p<0.001. **C**: Confocal images of F-actin (magenta) and cortical granule (CG Content; green) distribution in naturally fertilised WT (left), *par2a* mild (middle) and *par2a* severe (right) mutant eggs at 30 mpf. *par2a* mutants show retention of F-actin meshwork that correlates with retention of cortical granules at cortex. **D**: Quantification of CG content staining at cortex in fertilised WT, mild and severe *par2a* mutant eggs at 30 mpf. n=7, 3, 4; Mann-Whitney test; * = p<0.05; ** = p<0.01. **E**: Fluorescent eGFP superimposed on Nomarski image of naturally fertilised WT and mild *par2a* mutants injected with fluorescent latex beads at the vegetal pole (left column). Bead movement is recorded one hour later (right column). Arrow heads highlights latex bead position. **F**: Average distance travelled by injected fluorescent latex beads over 1 hour in naturally fertilised WT and mild *par2a* mutant eggs. n=5; Mann-Whitney test; ** = p<0.001. Scale bars: A, E = 200µm; C = 20µm.

Unlike mammals in which egg activation is initiated by sperm binding, egg activation in fish occurs upon spawning, can be generated experimentally by exposure to water, and does not require sperm (**Fig. 2A**). Indeed, attempts to activate the eggs of *par2a* females *in vitro* without sperm also failed, indicating that inability to undergo fertilisation *per se* is not the primary defect in eggs lacking *par2a* (**Supplementary Movie 3**).

### Cortical granule release and cytoplasmic streaming fail in *par2a* mutants

We quantified the hallmarks of egg activation in mild and severe clutches. The ratio of diameters of the chorion to egg body was significantly reduced in both mild and severe *par2a* mutants compared to WT, with severe mutants having chorion diameter only slightly larger than the egg body after 60 mins post fertilisation (**Fig. 2A, B**). Chorion elevation is driven by Cortical Granule Exocytosis (CGE), when cortical granules (CGs) docked at the inner surface of the cortex, fuse with the plasma membrane and release their contents into the perivitelline space. This inflates the chorion outwards as well as hardens it (3). The process is accompanied by cortical actin rearrangement of the egg surface within 5 minutes post egg activation (mpa), where the local network of F-actin is disassembled (3, 38). To determine if impaired chorion elevation in *par2a* mutants is due to defective CGE, we examined cortical actin distribution using AlexaFluor-546 Phalloidin. At 30 seconds after activation, WT eggs show a meshwork of F-actin under the plasma membrane. A similar meshwork is noted in activated eggs of both mild and severe *par2a* mutants (**Fig. S4E**). In WT this rapidly reorganises to become a smooth distribution of actin staining by 5 mpa. However, *par2a* mutant eggs fail to show this reorganisation and retain an intensely stained meshwork appearance (**Fig. S4E**). Even by 30 minutes post activation, fertilised mild and severe *par2a* eggs retained this cortical actin meshwork, whilst it was dissaembled in WTs (**Fig. 2C)**. As the actin meshwork is thought to act as a barrier to CG exocytosis, we directly stained CGs using FITC-conjugated *Maclura pomifera* Lectin (FITC-MPL). We observed that in contrast to WT which had almost complete release of CGs by 5 mpa, *par2a* mutants retained CGs at the egg surface even at 30 mpf (**Fig. 2C; Fig. S4F**). By comparing mean FITC-MPL intensity in WT and *par2a* mutants at 30 mpa, we saw a significant increase in retention of cortical granules in mild mutants, and a stronger retention in severe mutants (**Fig. 2D**). This correlates with the severity of chorion elevation defect observed between severe and mild *par2a* mutants. Thus, we conclude that cortical F-actin disassembly and CGE fails in *par2a* mutant eggs, leading to defective chorion elevation.

After egg activation initiation, cytoplasm in the egg separates away from the yolk and streams towards the animal pole, which eventually becomes the blastodisc (4). Because *par2a* mutants show defective blastodisc formation, we tested if cytoplasmic streaming is impaired by injecting fluorescent latex beads at the vegetal pole and tracking their movement over 1 hour. We observed that in naturally fertilised WT eggs, most of the beads migrated to the animal pole, sitting at the forming blastodisc, however in fertilised *par2a* mutant eggs, injected beads showed little displacement (**Fig. 2E; Supplementary Movie 4**). Quantification of latex bead displacement showed a significant reduction in *par2a* mutants compared to WT (**Fig. 2F; Fig. S4G**). Together, these results implicate a novel role of Par2a in egg activation and subsequent blastomere cleavage.

### Ca^++^ wave propagation is reduced in *par2a* mutants during egg activation and cell cleavage

To investigate if the defects observed in *par2a* mutants are Ca^++^ dependent as seen for serine protease inhibitors, we crossed the this mutant to the *Tg(actb2:GCaMP6s)*^*lkc2*^ transgenic line. Eggs from homozygous *par2a* mutant females were collected and GCaMP6s signals recorded by fluorescent time lapse imaging following E2 activation. This revealed that compared to WT, *par2a* mutant eggs showed attenuation of the Ca^++^ wave after activation. In all mutants, an initial focus of Ca^++^ was seen, but this showed reduced intensity and duration, and failed to propagate as extensively as in WT. Extent of attenuation correlated with severity of subsequent phenotype, where embryos with most reduction in chorion lift and blastodisc formation had the smallest Ca^++^ wave (**Fig. 3A-B; Supplementary Movie 5, 6**). Levels of calcium were reduced from the outset and remained so for the duration of imaging. Area under the curve analysis indicated significant attenuation of Ca^++^ signals for both severe and mild *par2a* mutants (**Fig. 3C**). Timelapse analysis suggested that in severely affected eggs, there was an initial puff of calcium, however this did not progress to a propagated wave (**Supplementary Movie 6**).

**Figure 3:**
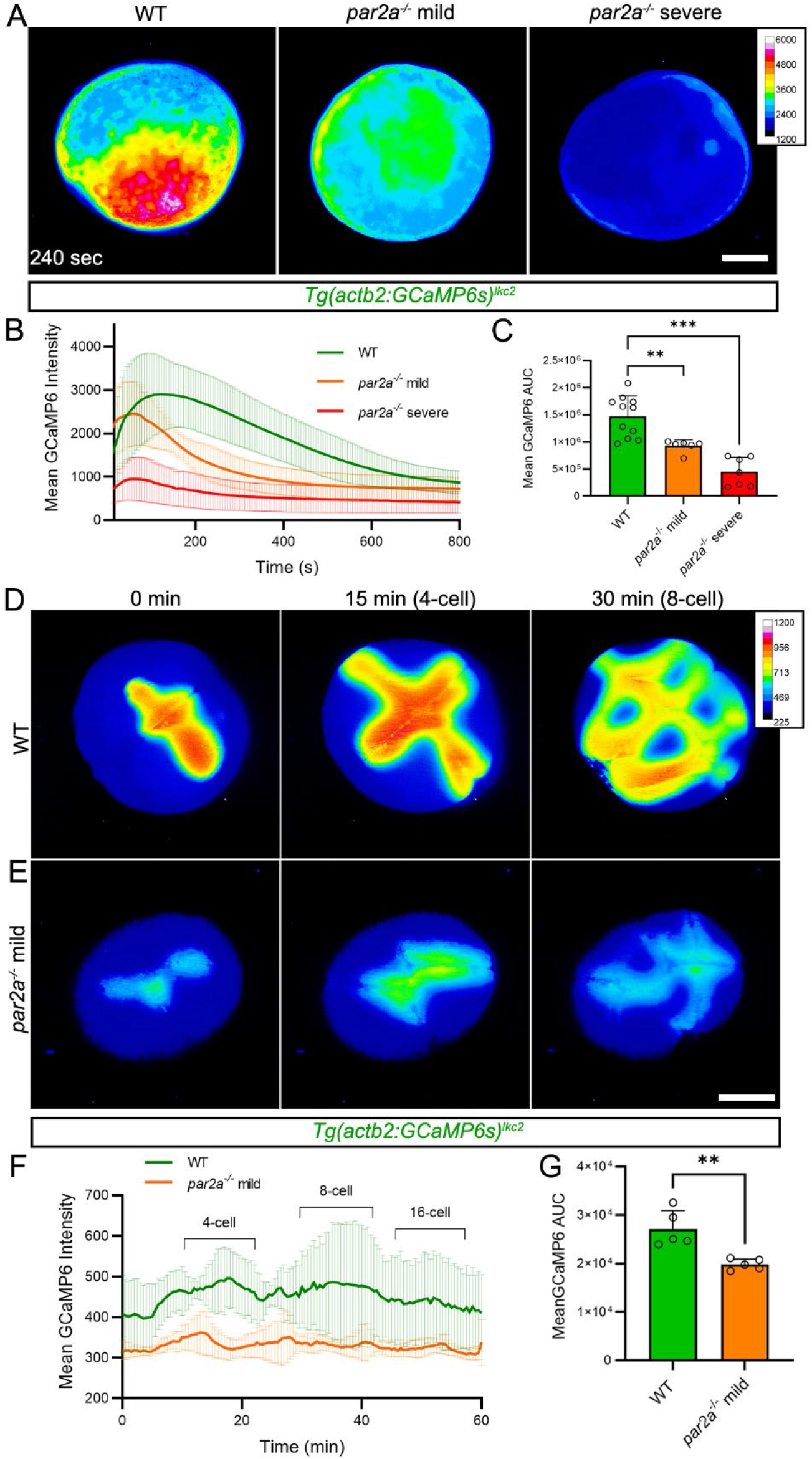
*par2a* mutants show defective Ca^++^ wave propagation during egg activation and blastomere cytokinesis. **A:** Fluorescent images of *Tg(actb2:GCaMP6s)*^*lkc2*^ indicating comparative Ca^++^ levels at 240 seconds post egg activation in WT, mild and severe *par2a* mutant eggs. **B:** Changes in GCamP6s intensity over 10 minutes from the *Tg(actb2:GCaMP6s)*^*lkc2*^ Ca^++^ reporter transgene of WT (green), mild *par2a* (yellow) and severe *par2a* (red) mutants following egg activation. **C:** Graph of corresponding Area Under Curve analysis of GCaMP6s levels presented in (B). n=11,6,7; Mann-Whitney test; ** = P<0.01; *** = p<0.001. **D-E:** Projected lightsheet images of *Tg(actb2:GCaMP6s)*^*lkc2*^ in naturally fertilised WT (D) and mild *par2a* mutant eggs (E) during cell division over 30 minutes starting from 2-cell stage. **F:** Changes in GCaMP6s intensity over 1 hour in WT (green) and mild *par2a* mutant embryos (yellow) during early blastomere division. Acquisition begins at 20 mpf and cell division events are indicated above WT by brackets. **G:** Graph of corresponding Area Under Curve analysis of GCaMP6s levels presented in (F). n=5; Mann-Whitney test; ** = p<0.01. Scale bars: A, E = 200µm.

As mildly affected *par2a* mutant eggs form blastomeres, we were able to examine if the defects in blastomere cytokinesis was associated with disruptions in normal Ca^++^ dynamics at the cleavage furrow. Characteristic, dynamic Ca^++^ waves at the furrow were apparent in WT, with transient slow calcium waves at the positioning of each furrow (initiation wave), which are then propagated as maturation waves along the furrow during deepening and zippering phases (**Fig. 3D; Supplementary Movie 7**) (17, 35). Although there were weak transients at the nascent furrows of *par2a* mutants (likely initiation waves), these failed to propagate effectively, failing to fully extend laterally across the division plane (**Fig. 3E; Supplementary Movie 7**). This coincided with loss of the orthogonal arrangement of cleavages in these embryos. Quantifying intensity of calcium transients during early blastomere cleavages showed a significant and sustained reduction in the mutants (**Fig. 3F-G**). These experiments indicate that *par2a* is required for intensity and propagation of calcium waves during egg activation and at cleavage furrows during division of the blastomeres.

### Defects in *par2a* mutants are rescued by increasing intracellular Ca^++^

To test if this loss of Ca^++^ indeed does account for the egg activation and cell division defects in *par2a* mutants, we asked if experimentally increasing intracellular Ca^++^ would rescue any aspects of the phenotype. The ionophore, Ionomycin, raises intracellular Ca^++^ in zebrafish eggs leading to activation (9). 2µM of ionomycin was added to eggs from a severely affected clutch of MZ*par2a* mutants immediately after natural spawning. After 1.5 hours, all untreated embryos showed lack of activation, with negligible chorion elevation and no overt blastodisc (**Fig. 4C**). However, individuals from the same clutch treated with ionomycin showed strong significant rescue of egg activation, including significant chorion elevation, prominent blastodisc and some blastomere cell division (**Fig. 4D, E**). However, these cell divisions were abnormal and treated *par2a* mutant embryos failed to successfully gastrulate. Treatment of WT with this concentration of ionomycin also resulted in blastomeres of unequal sizes (**Fig. 4A-B, E**), suggesting that Ca^++^ levels must be tuned appropriately for successful cleavage. A similar disruption of cleavage furrows by calcium ionophores has been noted in *Drosophila* spermatocytes (39).

**Figure 4:**
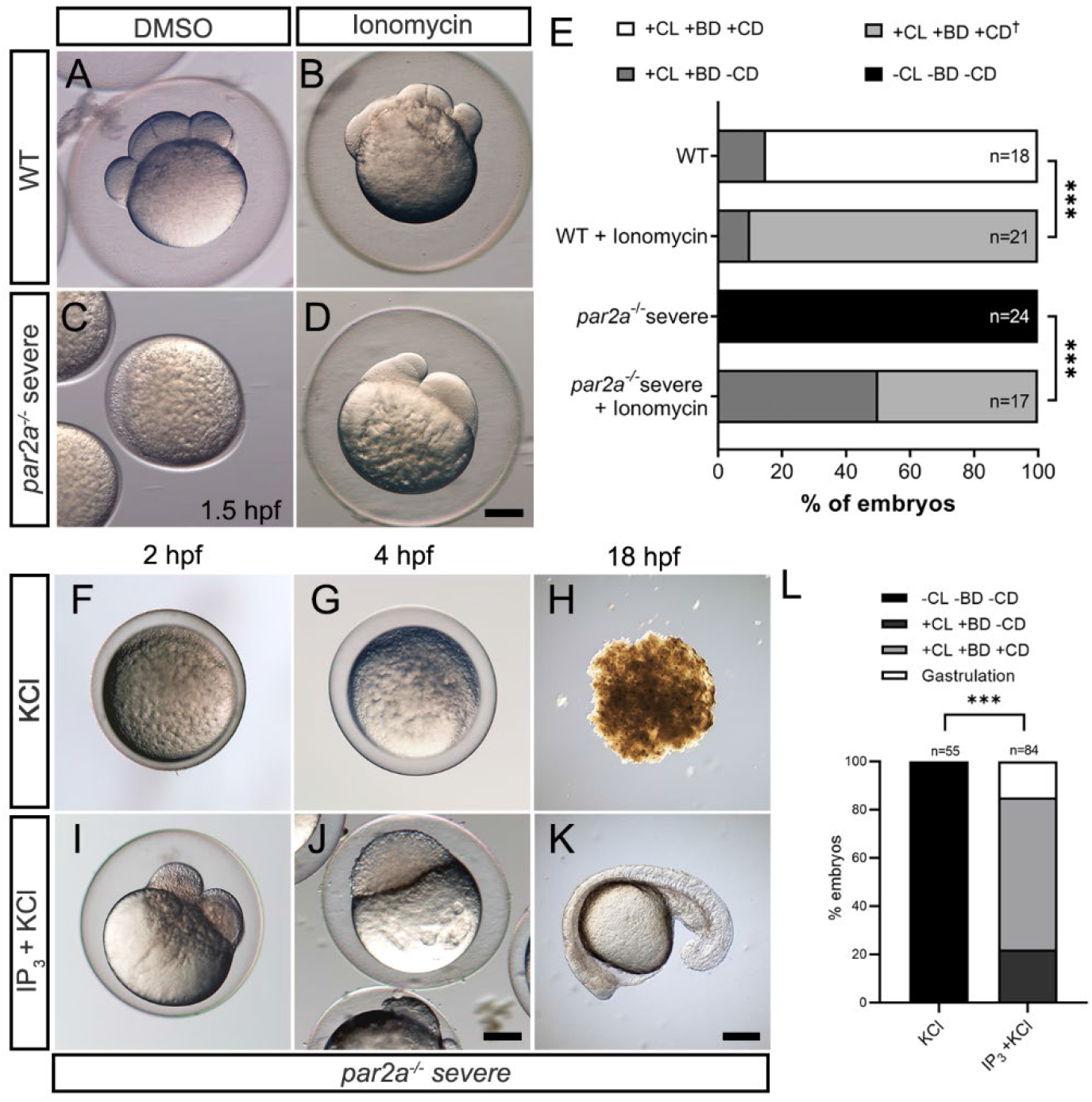
Increasing intracellular Ca^2+^ either by Ionomycin or IP_3_ rescues egg activation defects in *par2a* mutants. **A-D:** Images of WT (A, B) or severe *par2a* mutants (C, D) embryos derived from natural spawning and treated with 0.1% DMSO (A, C) or 2µM Ionomycin (B, D). **E:** Proportion of WT and severe *par2a*^*-/-*^embryos showing egg activation and blastomere division phenotypes at 1.5 hpf following treatment with 0.1% DMSO or 2µM Ionomycin. Key: CL: Chorion Lift, BD: Blastodisc, CD: Cell Division +: Present, -: absent. ^†^: Abnormal. Chi-squared analyses; *** = p<0.001. **F-K:** Images of naturally fertilised severe *par2a* mutant embryos injected with KCl alone (F, G, H) or with IP_3_ in KCl (I, J, K) at 2 hpf (F, I), 4 hpf (G, J) and 18 hpf (H, K). **L:** Proportion of egg activation and blastomere division phenotypes in fertilised severe *par2a* mutant eggs injected with IP_3_ in KCl or KCl alone. Key: CL: Chorion Lift, BD: Blastodisc, CD: Cell Division +: Present, -: absent. Chi-squared analyses; *** = p<0.001. Scale bars: D, J, K = 200µm.

### IP_3_ is necessary for zebrafish egg activation and is generated by Par2a activity

As outlined above, IP_3_ is sufficient to induce egg activation in a number of species including zebrafish (2, 40). IP_3_ sponges or IP_3_R inhibitors have demonstrated that IP_3_ signalling is also necessary for egg activation in sea urchins, *Xenopus*, and mammals (10, 41-43). Although Lee, Webb and Miller (22) have shown that zebrafish blastomere cleavage requires IP_3_R function, to our knowledge, necessity of IP_3_ earlier during zebrafish egg activation has not been directly tested. To do this we exposed WT eggs derived from natural spawning to increasing doses of the IP_3_R antagonist, 2-aminoethyl diphenyl borinate (2-APB). At lower concentrations, embryos successfully raised chorions and formed a blastodisc, whilst at 100µM, embryos failed to form a blastodisc, although they did show normal chorion elevation. Treatment with 500µM 2-APB abolished egg activation entirely, with no blastodisc, and no chorion elevation (**Fig S5A-E**). Thus, egg activation responds in a dose sensitive manner to IP_3_R inhibition and, as in other animal classes, IP_3_ is both necessary and sufficient for egg activation.

### Par2a regulates Ca^++^ during egg activation via IP_3_

This led us to determine if Par2a was regulating Ca^++^ in the egg via IP_3_. Firstly, we tested the sensitivity of mildly affected *par2a* mutants to low levels of 2-APB, reasoning that mildly reducing IP_3_R activity would only synergise with mild *par2a* mutants if the latter was also disrupting IP_3_ signalling. Mild *par2a* mutants partially raised a chorion and had a limited blastodisc but aborted blastomere cytokinesis (**Fig. S5H, J**). However, treatment with 50µM 2-APB significantly enhanced the severity of the egg activation phenotype of mildly affected *par2a* mutants, with all embryos completely lacking a blastodisc and failing to raise a chorion at all, and in contrast to the mild effects of activation this concentration of 2-APB had on WT embryos (**Fig. S5F, G, I, J**). Thus, *par2a* mutants are acutely sensitive to mild reduction of IP_3_R activity.

We also tested if supplementation of exogenous IP_3_ ligand can rescue *par2a* defects as observed for serine protease inhibitors. Injection of 20pmol IP_3_ into naturally fertilised clutches of both mildly and severely affected *par2a* mutants was able to rescue egg activation defects with high efficiency. In clutches derived from severe *par2a* females, injection of IP_3_ led to all embryos raising a chorion and forming a blastodisc, whereas this never occurred in KCl injected siblings (**Fig. 4F, I, L**). Many of these rescued embryos were able to undergo normal blastomere division and remarkably many could gastrulate to form a body axis at 18hpf whereas all uninjected siblings failed to even generate a blastodisc and underwent necrosis by 18hpf (**Fig. 4G-H, J-L**). Rescue of mildly affected *par2a* mutant clutches was even more pronounced, with almost 40% of embryos completing gastrulation, forming a normal body axis and surviving past 24 hpf compared to KCl injected controls (**S6A-C**). We conclude that *par2a* regulates IP_3_ levels to ensure effective egg activation.

### Supplementing IP_3_ restores cortical granule exocytosis and calcium dynamics in *par2a* mutants

Provision of IP_3_ rescued chorion elevation in severe *par2a* mutants, hence we checked if this was due to restoration of CGE by examining cortical F-actin distribution and CG content using Phalloidin and FITC-MPL stain, respectively. KCl injected severe *par2a* mutant embryos showed clusters of circular cortical F-actin staining which corresponded with high numbers of non-exocytosed CGs located at the cortex (**Fig. 5A**). By contrast, IP_3_ injected mutants displayed smooth and even cortical F-actin distribution with significantly reduced number of CGs indicating occurrence of CGE (**Fig. 5B-C**). To confirm if exogenous IP_3_ restores intracellular Ca^++^ in *par2a* mutants, we compared *in vivo* Ca^++^ dynamics using the T*g(actb2:GCaMP6s)*^*lkc2*^ reporter line in IP_3_ and KCl injected severe *par2a* mutants. Whilst KCl injected controls showed only a small brief increase in Ca^++^ transient, IP_3_ injected *par2a* mutant embryos showed a significant spatial increase in Ca^++^ levels with a longer wave duration (**Fig. 5D-E; Supplementary Movie 8**).

**Figure 5.**
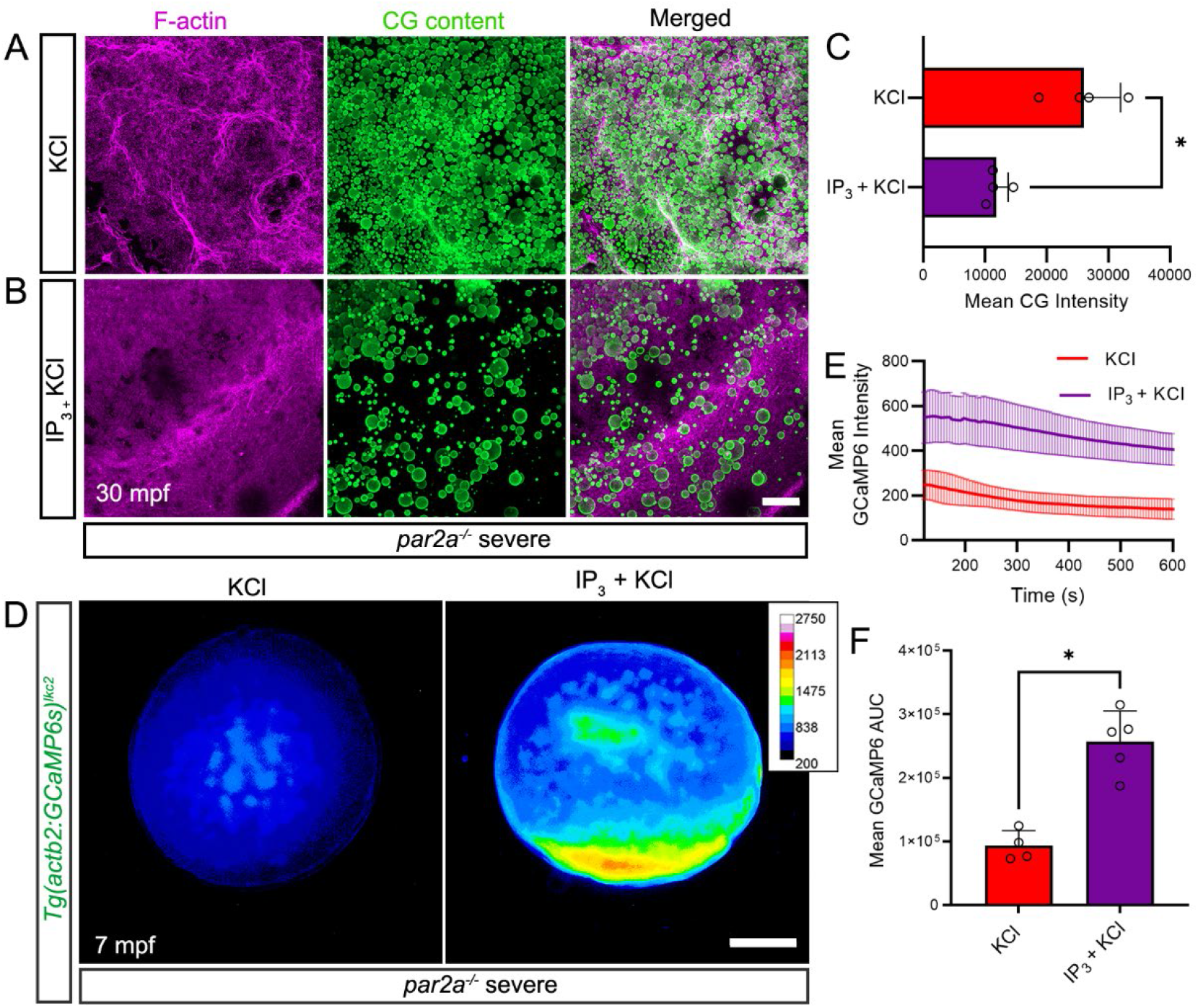
IP_3_ rescues egg activation defects by increasing intracellular calcium spatially. **A-B:** Confocal images of F-actin (left; magenta), and cortical granule (middle; CG Content; green) distribution at the cortex of naturally fertilised severe *par2a* mutant eggs at 30 mpf. Eggs were injected with KCl alone (A) or with 20pmol IP_3_ in KCl (B). **C:** Quantification of CG content staining intensity in fertilised severe *par2a* mutant eggs injected with KCl alone (red bar) or with IP_3_ in KCl (purple bar) n=4; Mann-Whitney test; * = p<0.05 **D:** Fluorescent images of *Tg(actb2:GCaMP6s)*^*lkc2*^ indicating calcium levels of naturally fertilised severe *par2a* mutant eggs at 7 minutes following injection with KCl alone (left) or with IP_3_ in KCl (right). **E:** Changes in GCaMP6s intensity reported by the *Tg(actb2:GCaMP6s)*^*lkc2*^ transgene over 10 minutes in fertilised severe *par2a* mutants eggs following injection with KCl alone (red line) or with IP_3_ in KCl (purple line). **F:** Graph of corresponding Area Under Curve analysis of GCaMP6s levels presented in (E). n=4, 5; Mann-Whitney test; * = p<0.05. Scale bars B = 50µm; D = 200µm.

## Discussion

We have identified the first cell surface receptor directly required for Ca^++^ regulation during egg activation. Our *par2a* maternal zygotic CRISPR mutants showed a strong egg activation defect leading to failure to raise a chorion and form a blastodisc. This was also evident in non-fertilised eggs, indicating this defect was not a fertilisation block, rather disrupting egg activation specifically.

As a GPCR activated by extracellular serine protease activity, implication of a Par2 paralogue in egg activation unifies a number of disparate observations. Firstly, trypsin inhibitors block egg activation in fish species, including, as demonstrated here, in zebrafish (8). We show that this is due to a significant and sustained reduction in Ca^++^ waves in the egg. Complimenting this, addition of exogenous trypsin to Xenopus oocytes was previously demonstrated to liberate Ca^++^ from internal stores (44). It was shown subsequently that this activity is mediated by the xenopus Gq and G_14_ homologues (45). Gq is also required for maximal Ca^++^ release in the sea urchin (46). Par2 has been implicated in activation of Gq in both mouse and zebrafish (31-33), and experiments in xenopus did indeed propose that the effects of trypsin on egg Ca^++^ was mediated by a trypsin receptor, even suggesting PAR-2 as the potential receptor (47). Our injection experiments indicate that supplementation of IP_3_ rescues both trypsin inhibition and loss of maternal Par2a, implying that a PLC isozyme lies downstream.

So far, two Phospholipase C isozymes have been implicated in egg activation. PLCζ is transferred directly into the egg from the sperm upon fertilisation in mammals, whilst PLCγ is important in egg activation of non-mammalian species and is activated by cytosolic kinases such as those of the FAK and Src families (5). The external stimulus activating these kinases is unclear. As Gq is known activator PLCβ, it is tempting to implicate this isozyme downstream of Par2a in the generation of IP_3,_ and thus Ca^++^ release, during zebrafish egg activation. Egg derived PLCβ has been shown to be required for maximum amplitude of Ca^++^ transients in mouse eggs (48). This is similar to our observations of Par2a mutants, which show a reduced intensity of Ca^++^ waves. In all movies, even in strong mutants, we noted presence of a muted and brief calcium wave, which failed to propagate effectively. This would suggest that PLCβ may have a specific role in Ca^++^ wave amplification and propagation in zebrafish, rather than initiation, although its involvement will require direct testing. Similarly, experiments in xenopus have also indicated that PLC activation might occur through a non-canonical mechanism, independent of heterotrimeric G-proteins (43, 49). Definitively positioning of Gq and PLCβ as downstream mediators of Par2a in zebrafish egg activation remains an important goal.

We found that the same *par2a* mutation had vastly different expressivity in eggs from different females. While there is little variation within a clutch, or between clutches from the same female, we saw that some females consistently gave MZ*par2a* clutches that were able to form a blastodisc and undergo limited, abnormal, blastomere cleavage. This allowed us to identify a later function for Par2a in regulating calcium dynamics at the blastomere cleavage furrow. No MZ*par2a* mutant ever successfully gastrulated. The reason for this variable expressivity is unknown. We hypothesised that compensation from maternal *par2b* might account for this variation, however some MZ*par2a; par2b* double mutant females also gave mildly affected clutches. There may be compensation from another Protease activated receptor homologue in certain females. Alternatively, these eggs might have upregulated parallel IP_3_ or Ca^++^ regulating pathways, such as Src family kinase activation of PLCγ.

At the blastomere cleavage furrow, Ca^++^ is released in distinct waves corresponding to furrow positioning, propagation, and deepening (20). Its precise functions are not fully understood although there is strong evidence it is required for cytoskeleton reorganisation and vesicle exocytosis during apposition (20). Analysis of the *nebel* mutant which has defective slow calcium waves has suggested that Ca^++^ acts to propagate and deepen furrows through a Calmodulin and CamK mechanism (23). The regulation of calcium at the furrow is unclear, with both ER and mitochondria sources available (23). Previous work had indicated that IP_3_ is essential for Ca^++^ release during blastomere cleavage furrow deepening (22), although how this is generated at this location is not known. Our work shows that in mild *MZpar2a* mutants, Ca^++^ transients at the incipient furrows are not sustained or propagated, leading to aberrant cytokinesis and mis-sized daughter cells. This identifies Par2a as an essential regulator of IP_3_ levels at the zebrafish blastomere cleavage furrow, and acts with a known role of Store Operated Calcium Entry to replenish ER Ca^++^ levels (21). In mammalian cells, the localisation of PLCβ1 at the cleavage furrow also suggests a mechanism through Gq to regulate IP_3_ levels (50), although there is no definitive identification of Par2 orthologues in mammalian blastocysts or oocytes.

More generally, to what extent the role of Par2 in egg activation, and early embryo cell cleavage is conserved in mammals is not yet clear. Immunostaining hinted that Par2 is expressed on mouse oocytes and can be activated by the sperm derived protease, Acrosin (51), although there is no evidence from Par2 knockout studies that it is necessary for mouse egg activation. Prior to mammalian blastocyst implantation, the endometrium senses the presence of the invading embryo and initiates a number of intracellular signals, including waves of Ca^++^. This has recently been shown to be mediated by endometrial expression of PAR2, which is activated by trypsin derived from the invading blastocyst, and which acts through PLC and IP_3_R, as seen in zebrafish egg activation (52). Whether PAR2 or related receptors also plays role in mammalian oocyte maturation itself or in blastocyte cell division, as seen in zebrafish, requires testing.

As egg activation can occur without fertilisation and requires Par2a, then the protease activating Par2a must be expressed in the egg as well. It is critical to identify this protease and moreover, determine how it is only activated upon spawning. One possible mechanism is simply a dilution away of serine protease inhibitors, known to be expressed in fish ovaries (7). Similarly, release into water could alter the pH or ionic conditions that activate the protease. Alternatively, the protease could be sequestered in cortical granules and only released upon initial activation, where it can then cleave the extracellular domain of Par2a. This may then lead to a wave of Ca^++^ as Par2a activation spreads around the cortex and granules are progressively released, acting as a form of Calcium Indiced Calcium Release.

## Materials and Methods

### Zebrafish maintenance and lines

Zebrafish lines were raised and maintained in NTU zebrafish facility under standard protocols (28^°^C; 14hr-light/10hr-dark cycle) in compliance with guidelines provided by National Advisory Committee for Laboratory Animal Research. *In-vivo* Ca^++^ imaging was performed using the *Tg(actb2:GCaMP6s, myl7:mCherry)*^*lkc2*^ transgenic line (termed *Tg(actb2:GCaMP6s)*^*lkc2*^ hereafter) (32, 35). All experiments were performed under IACUC number #A18002.

### Egg collection

Embryos collected through natural mating were incubated at 28°C in E3 embryo medium (5mM NaCl, 0.17mM KCl, 0.33mM CaCl_2_, 0.33mM MgSO_4_). Un-activated/un-fertilized eggs were collected by squeezing eggs from female fish directly as per methods adapted from (53) following anaesthesia with 0.02% Tricaine in E3 (MS-222 Sigma buffered to pH 7.0).

### *In-vitro* egg activation and fertilisation

For experiments that did not require fertilisation, eggs were covered in buffered Hank’s saline (0.137M NaCl, 5.4mM KCl, 0.25mM Na_2_HPO_4_, 1.3mM CaCl_2_, 1.0mM MgSO_4_, 4.2mM NaHCO_3_) containing 0.5% Bovine Serum Albumin (BSA) to prevent egg activation (34). To initiate egg activation, 0.5% BSA was diluted out with 0.5x E2 medium (7.5mM NaCl, 0.25mM KCl, 0.5mM MgSO_4_, 75μM KH_2_PO_4_, 25μM Na_2_HPO_4_, 0.5M CaCl_2_, 0.35mM NaHCO_3_; buffered to pH 7.0 with HCl). For *in vitro* fertilisation, dissected male testes were macerated in ice-cold buffered Hank’s saline with 0.5% BSA. Eggs were fertilized *in vitro* by adding sperm suspension mixture to eggs, and then released for development by transferring to 0.5x E2 embryo medium. Staging of embryos were as per (54) while embryos prior to 1-cell stage were charted according to time from activation/fertilisation.

### Generation of *par2* knockouts

Mutations in the *par2a* and *par2b* genes were generated by CRISPR/Cas9 mutagenesis with single guide RNA (sgRNA) targeting exon 2 of each paralogue at a site corresponding with the 2^nd^ extracellular loop of the transmembrane helix determined by protein topology prediction provided by TMHMM v2.0 server (55). Oligo templates for sgRNA transcription containing T7 promoter binding site was designed according to (56) utilising the following crRNA and PAM (uppercase) sequence (*par2a -* 5’ gggtcgagtgacatcatggcAGG 3’; *par2b -* 5’ ggagatgtgcaaagtatcagTGG 3’). The completed sgRNA template was obtained through PCR using TaKaRa PrimeSTAR® Max DNA polymerase and transcribed with MEGAshortscript™ T7 Kit from Invitrogen. *Cas9* RNA was transcribed from the pCS2-nCas9n plasmid (57) using the mMESSAGE mMACHINE™ SP6 Transcription Kit (Invitrogen). sgRNA and Cas9 RNA were diluted in 1x Danieau’s buffer (5mM HEPES (pH 7.6), 58mM NaCl, 700µM KCl, 400µM MgSO_4_.7H_2_O, 600µM Ca(NO_3_)_2_) with Phenol Red and injected into embryos at the 1-cell stage. A sample of injected embryos were sequenced at 24 hpf to evaluate mutagenesis efficacy and remaining embryos were raised to adulthood. To identify founders, adults were incrossed and progeny were sequenced for mutations. Double mutants (*par2a*,*b*) were generated by injecting *par2a* sgRNA into the established *par2b* mutant line.

### Fluorescent labelling

*In vitro* or naturally fertilised eggs were fixed at respective timepoints in 4% paraformaldehyde in PBS at 4°C overnight. For visualizing cortical granules, fixed embryos were washed in PBST (0.1% Tween^TM^ 20 in PBS), permeabilized in 0.2% Triton-X in PBS for 2 hours, dechorionated and labelled by incubation overnight at 4ºC in PBST with 50µg/ml FITC-conjugated *Maclura pomifera* Lectin (F-3901-1, EY Laboratories, Inc). To examine cortical F-actin distribution, embryos were incubated overnight at 4ºC in PBST containing a 1:500 dilution of Alexa Fluor™-546 Phalloidin (A12379, Invitrogen™). Where possible, FITC-MPL and Phalloidin stainings were performed simultaneously. Embryos were subsequently washed in PBST and stored in glycerol for imaging.

### Tracking Cytoplasmic Streaming

Embryos collected from natural spawning were injected at the vegetal pole at 5 mpf with 1nl of 1:5 fluorescent yellow-green carboxylate-modified polystyrene latex beads (L4530, Sigma-Aldrich^®^) diluted in H_2_O. Embryos were then immediately mounted in 0.8% Low Melt Agarose (LMA) (Mo Bio Laboratories) in 0.5x E2 medium and orientated laterally on 15 mm glass-bottom imaging dishes (NEST^®^ Biotechnology, 801002) for imaging. Distance of travel was measured by taking two images on an upright Zeiss AxioImager M2 microscope at 0mins and 60mins after mounting. Timelapse movies of bead movement were taken using an upright Zeiss LSM800 confocal microscope with a 45s interval time.

### Inhibitor and rescue treatments

For inhibitor treatments of eggs, squeezed eggs were held in Hank’s saline containing 0.5% BSA, sperm solution added and subsequently released for egg activation and fertilisation by washing with 0.5x E2 embryo medium containing inhibitor solution. To inhibit IP_3_R, 2-APB (D9754, Sigma-Aldrich^®^) was dissolved in Methanol and used at 50µM, 100µM or 500µM. 0.2% Methanol was used as control.

For protease inhibitor treatment, Aprotinin (14716, Cayman Chemical) and Benzamidine Hydrochloride Monohydrate (Benzamidine HCl) (B-050, Gold Biotechnology^®^) were dissolved in H_2_O. and diluted in 0.5x E2 embryo medium before adding to suspended egg sperm mix. Aprotinin and Benzamidine HCl were used at 5mg/ml and 44mg/ml respectively.

For Ionomycin treatment, naturally fertilised eggs were incubated in 0.5x E2 supplemented with 2µM Ionomycin (I0634, Sigma; dissolved in DMSO) with 0.1%DMSO as control. For IP_3_ rescue, 2nl of a 10mM IP_3_ solution (D-myo-Inositol 1,4,5-tris-phosphate trisodium salt; I9766, Sigma-Aldrich; dissolved in 0.7mM KCl) was injected into the yolk of naturally fertilised eggs. Injected eggs were observed at different time points as stated. For control comparison, eggs were injected with 0.7mM KCl solution only.

### Microscopy

Images were taken on an upright Zeiss AxioImager M2 compound microscope, an upright Zeiss LSM800 confocal microscope, a Zeiss AxioZoom V16 microscope and a Zeiss Light-sheet Z.1 microscope. Images were processed on Zen (ver. 2.3 Lite) or using Fiji (ImageJ, v1.53t). Embryos and eggs were mounted laterally in 0.8% LMA in 0.5x E2 medium in 15mm glass-bottom dishes (Nest Scientific).

Live low power images of embryos in E2 medium were taken using a Zeiss AxioZoom V16 microscope. Nomarski imaging was performed on the Zeiss AxioImager M2 microscope.

For visualizing early calcium dynamics, 0.5x E2 activated unfertilised eggs squeezed from *Tg(actb2:GCaMP6s)*^*lkc2*^ females or IP_3_ injected eggs were imaged directly on a Zeiss AxioZoom V16 microscope without mounting. For visualizing calcium dynamics at the cleavage plane, *Tg(actb2:GCaMP6s)*^*lkc2*^ embryos, collected by natural spawning, were dechorionated manually and mounted in 1% LMA in a 50µl volume capillary (BRAND™ 701908). Fluorescent timelapse series was then captured on a Zeiss Light-sheet Z.1 microscope.

### Image processing and quantification of GCaMP6 fluorescence

Mean intensities of projected fluorescent images were analysed using the Average Intensity function in Fiji. For fluorescent time-lapse images, an average intensity plot across time was generated by measuring the mean fluorescence corrected for background fluorescence, for each time frame. For each frame, a region of interest corresponding to the embryo outline was generated by a Create Selection function in Fiji.

### Statistical Analysis

Graphpad Prism software was used for statistical analysis. All graphs represent mean with standard deviation. Statistical tests used were Student’s t-test, Mann-Whitney test, Area Under Curve with Mann-Whitney test, and Chi-squared test, as stated in figure legends. P values were presented as * = p<0.05, ** = p<0.01 or *** = p<0.001, with 0.05 set as alpha in all.

## Notes

### Competing Interest Statement

The authors have declared no competing interest.

