## Supplementary Data for "Protease-activated receptor 2 links protease activity with calcium waves during egg activation and blastomere cleavage"

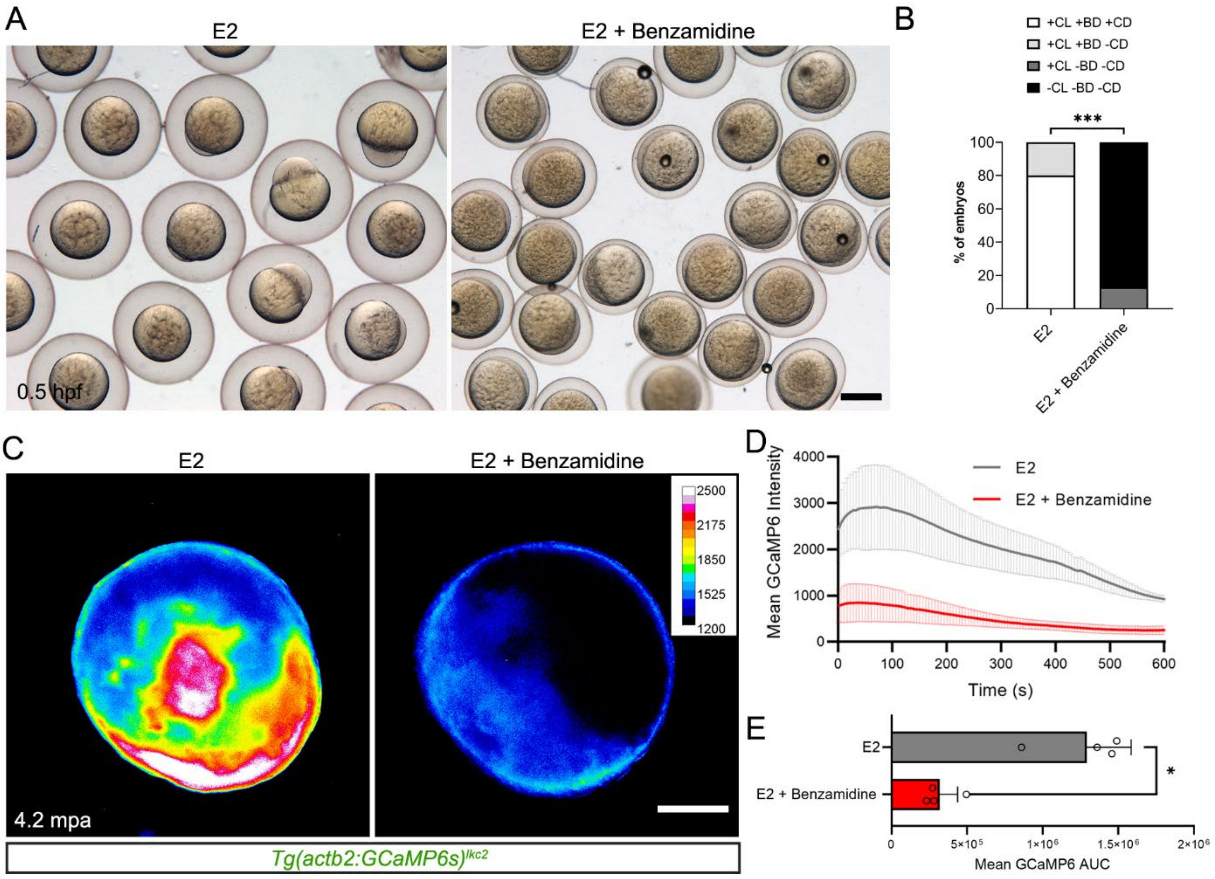


**Supplementary Figure 1: The serine protease inhibitor Benzamidine aborts egg activation and reduces $Ca^{2+}$ wave propagation.**

**A:** Eggs fertilised *in vitro* in E2 medium with or without 40mg/ml Benzamidine HCl. **B:** Proportion of embryos showing egg activation and blastomere division phenotypes for 40mg/ml Benzamidine HCl treatment vs E2 alone. Key: CL: Chorion Lift, BD: Blastodisc, CD: Cell Division +: Present, -: absent. Chi-squared analyses; *** = p<0.001; n = 100. **C:** Fluorescent images of unfertilised *Tg(actb2:GCamP6s)*$^{lkc2}$ eggs indicating $Ca^{2+}$ dynamics at 250 sec during egg activation in E2 or Benzamidine HCl. **D:** Quantification of changes in mean GCamP6s intensity from fluorescent timelapse of *Tg(actb2:GCamP6s)*$^{lkc2}$ eggs activated in E2 (grey line) vs 40mg/ml Benzamidine HCl treatment (red line) **E:** Corresponding statistical analyses of GCamP6s intensity AUC from (D). n=4; Mann-Whitney test; * = p<0.05. Scale bars: A = 500µm, C = 200µm.

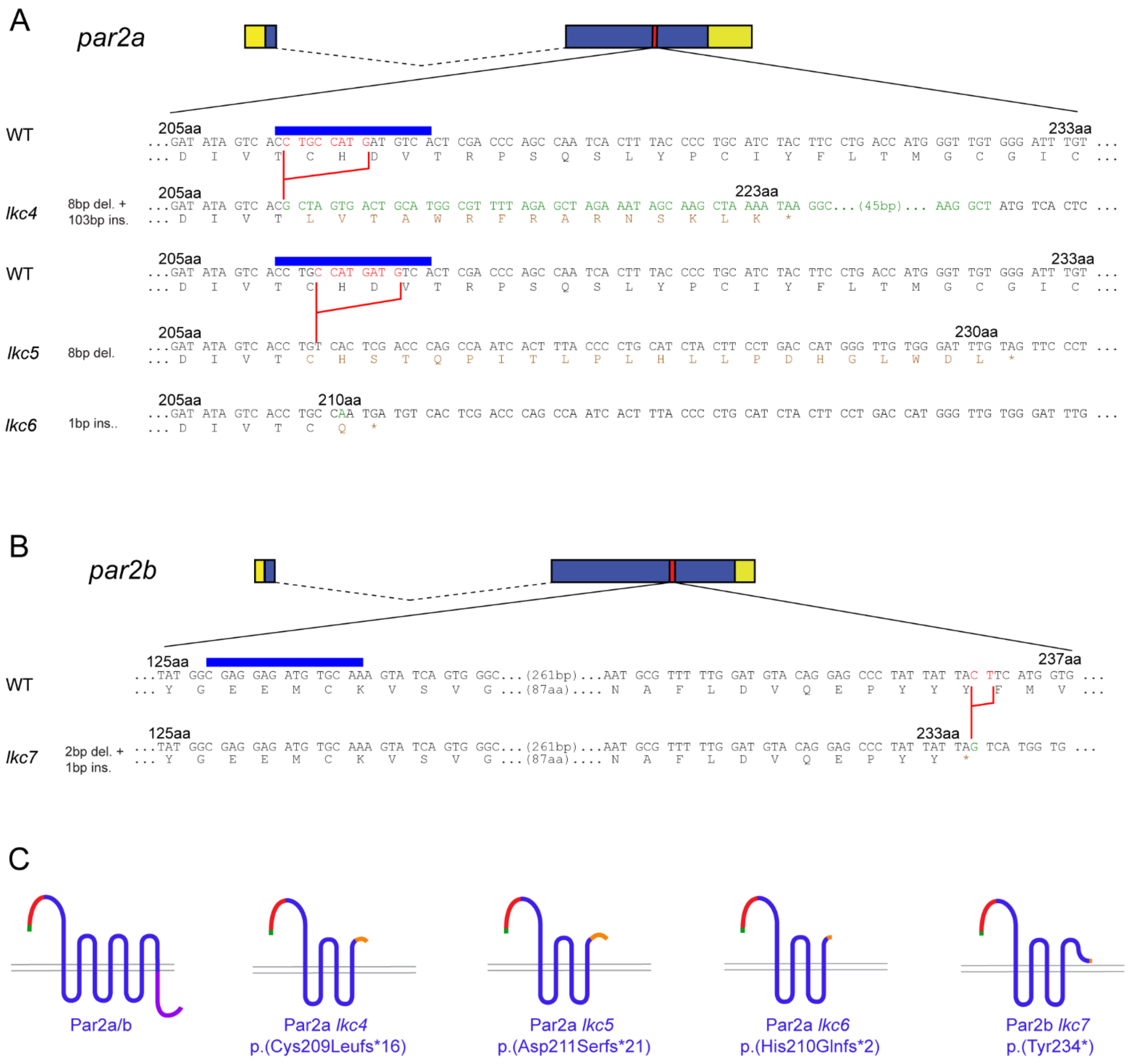


**Supplementary Figure 2: Generation of *par2a* and *par2b* mutant alleles**

**A - B:** CRISPR mutagenesis of *par2a* (A) and *par2b* (B). Intron-Exon structures are given above. Boxes correspond to exons and dashed lines represent introns. Both genes have 2 exons. Dark blue and yellow boxes represent coding and untranslated regions respectively. Red bars indicate approximate location of CRISPR target site with sequence given below for each allele (name and mutation summary given on left), with corresponding WT sequence for comparison. Blue bars designate CRISPR binding sites. Red and green text indicate deleted and inserted nucleotides respectively. Brown amino acid sequences indicates novel amino acids introduced by frame shift. **C:** Protein schematics for all Par2a and Par2b CRISPR alleles. Green, red, blue, purple and orange sections indicate signal peptide, tethered inhibition domain, 7-pass transmembrane region, intracellular tail and induced frame shift regions respectively. Corresponding allele names and summary given below.

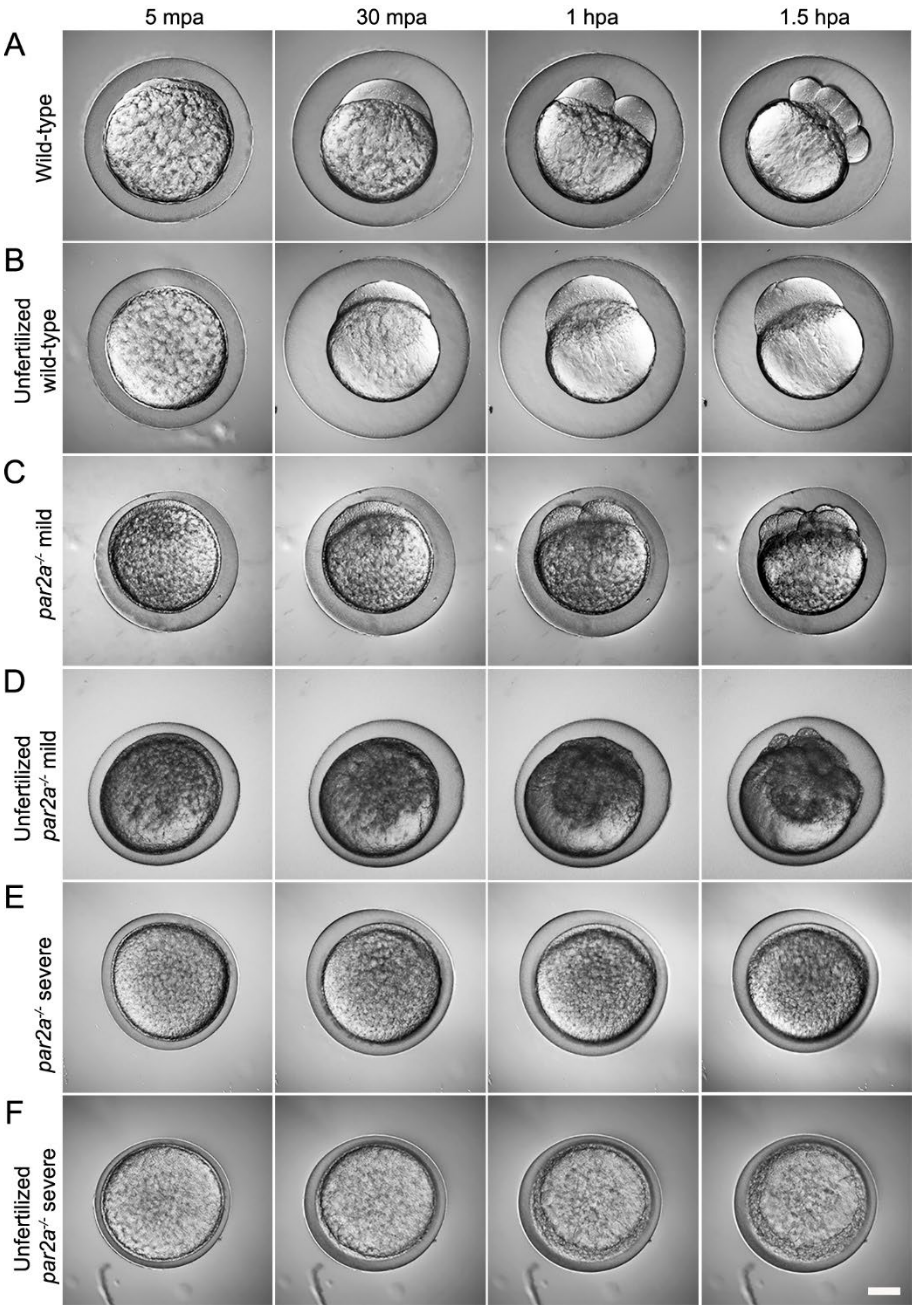


**Supplementary Figure 3: Aborted egg activation and cell division in *par2a* mutants**

Nomarski images of naturally fertilized (A, C, E) and unfertilised (B, D, F) eggs derived from WT (A, B), mild (C, D) and severe (E, F) *par2a* mutant females. Eggs were fertilised with WT sperm in A, C, E. Images were taken at 5 minutes (5 mpa), 30 minutes (30 mpa), 1 hour (1 hpa) and 1.5 hours post activation (1.5 hpa). Chorion elevation is reduced in the *par2a* mutant derived eggs (C-F) and a defective blastodisc forms only in the mild *par2a* mutant eggs (C, D), with none forming in either the fertilised or unfertilised severe mutant eggs (E, F). Scale bar: F = 200μm

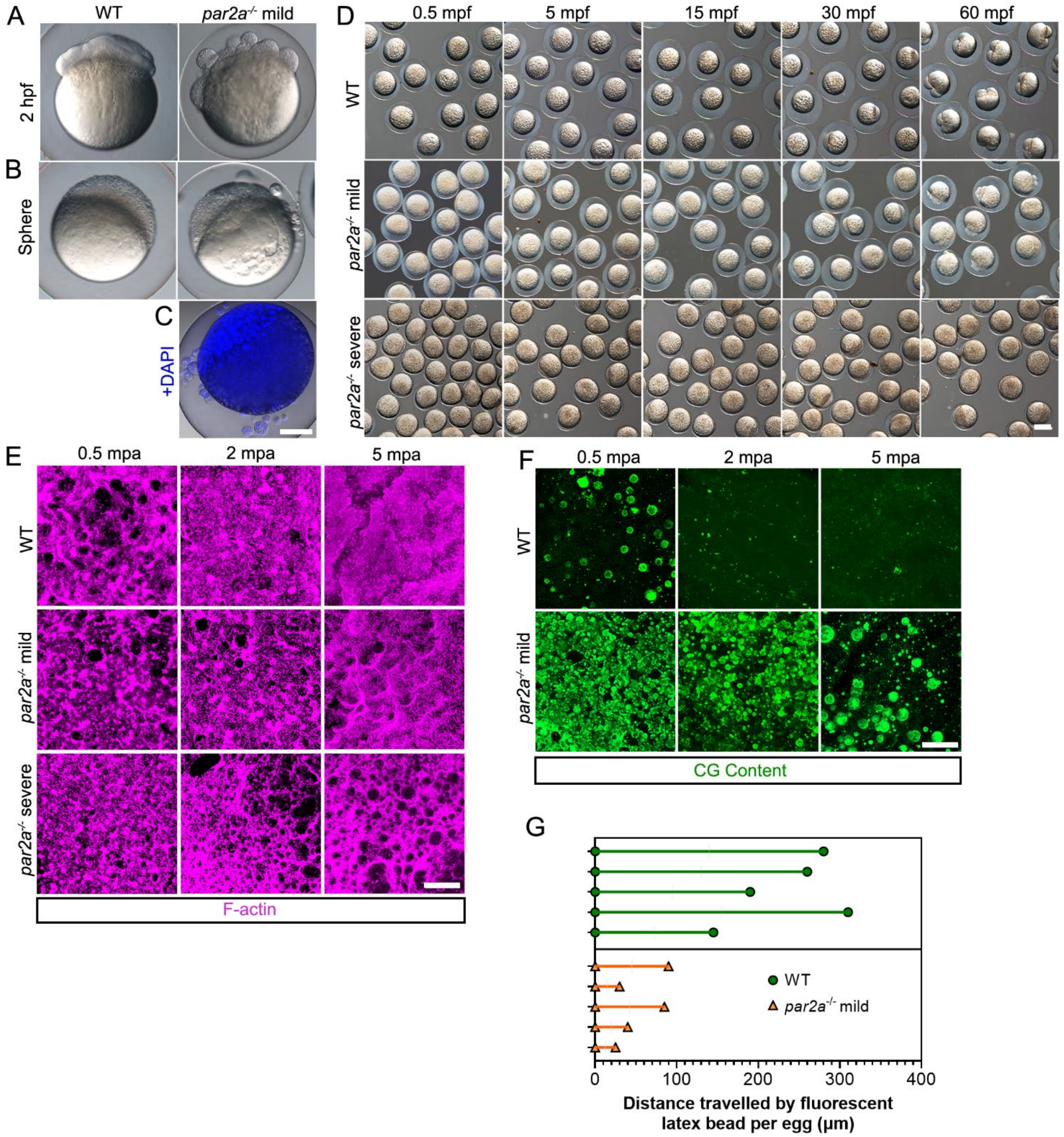


**Supplementary Figure 4: Loss of *par2a* generates egg activation defects.**

**A-C:** Nomarski images of WT (left) and mild *par2a*$^{-/-}$ mutants (right) derived from natural crosses at 2 hpf (A) and sphere stage (4 hpf; B). **C:** DAPI staining superimposed on Nomarski images of mild *par2a*$^{-/-}$ mutants at sphere stage **D:** Clutches of embryos from WT, mild and severe *par2a*$^{-/-}$ mutants generated by IVF and imaged at 0.5 mpf, 5 mpf, 15 mpf, 30 mpf and 60 mpf. **E:** Projected confocal images showing dynamics of cortical F-actin stained by AlexaFluor-546 Phalloidin at 0.5 mpa, 2 mpa and 5 mpa, in E2 activated WT (top), *par2a*$^{-/-}$ mild (middle) and severe *par2a*$^{-/-}$ mutant eggs (bottom). **F:** Projected confocal images showing dynamics of Cortical Granule release stained by FITC-MPL at 0.5 mpa, 2 mpa and 5 mpa, in E2 activated WT (top), and *par2a*$^{-/-}$ mild mutants (bottom) **G:** Distance travelled by injected fluorescent latex beads in individual fertilised eggs of WT (top; green) vs mild *par2a*$^{-/-}$ mutants (bottom; yellow) from start point (left) to end position at 1 hour. Each line represents the distance travelled by tracked beads in each embryo. Scale bars: C = 200µm; D = 500µm; E, F = 50µm.

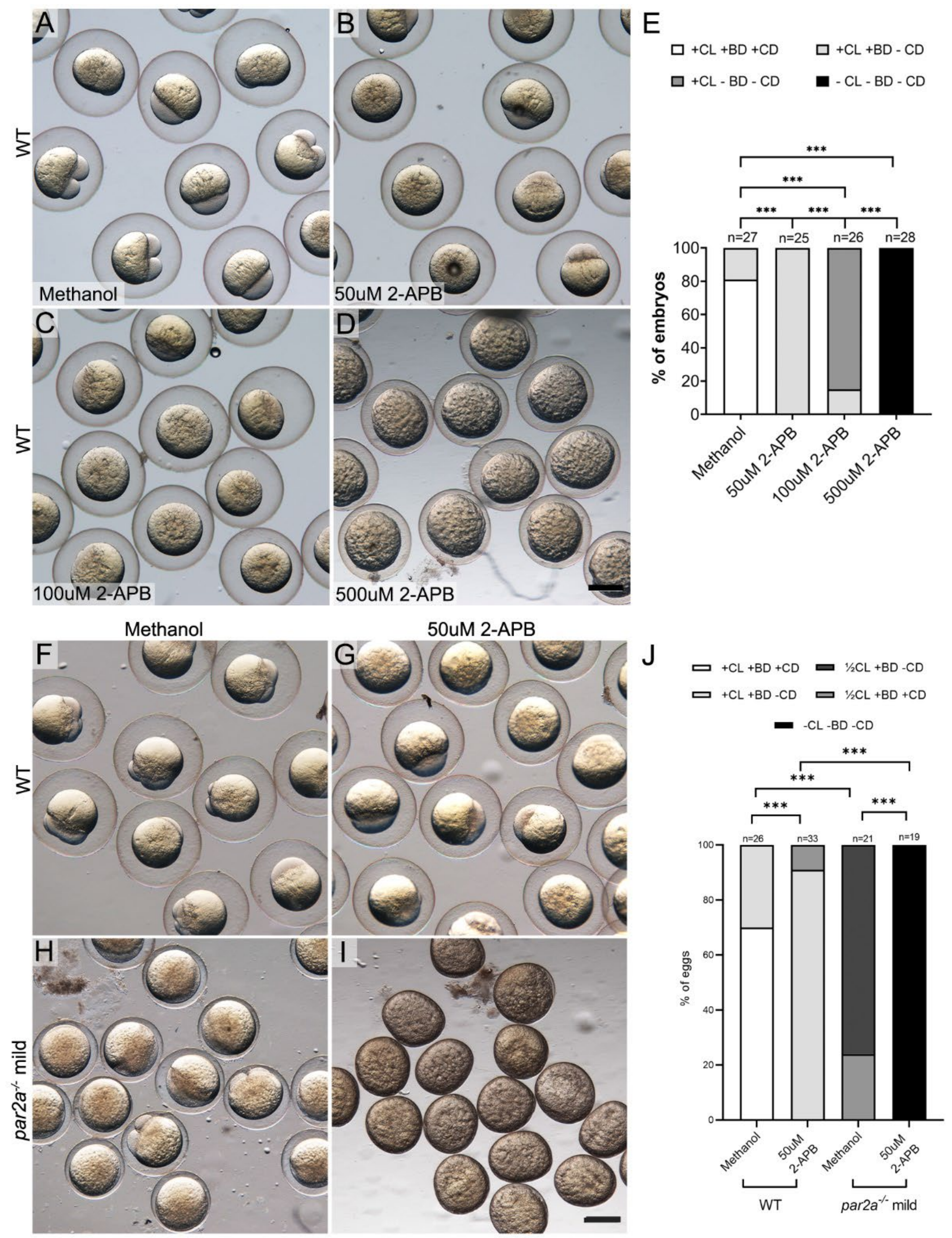


### Supplementary Figure 5: Egg activation of *par2a* mutants is highly sensitive to $IP_3R$ antagonism

**A-D:** WT embryos treated with methanol carrier (A), or 50µM (B), 100µM (C), and 500µM (D) of $IP_3R$ inhibitor, 2-APB. **E:** Proportion of egg activation phenotypes presented by different concentrations of 2-APB. Key: CL: Chorion Lift, BD: Blastodisc, CD: Cell Division +: Present, -: absent. Chi-squared analysis; *** = p<0.001. **F-I:** Naturally fertilised WT (F, G) and mild *par2a* mutant (H, I) embryos treated with methanol (F, H) or a low dose (50µM) of 2-APB. Low dose of 2-APB strongly exacerbates egg activation defects in mild *par2a* mutants. **J**: Counts of proportion of embryos in WT and mild *par2a* mutants showing extent of phenotype in low dose of 2-APB. Key: CL: Chorion Lift, BD: Blastodisc, CD: Cell Division +: Present, -: absent, ½: half reduced; Chi-squared analysis; *** = p<0.001. Scale bars: D, I = 500µm

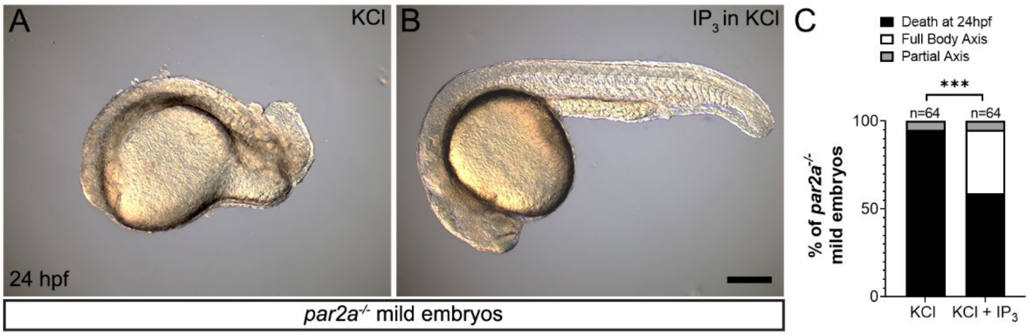


**Supplementary Figure 6: Injection of $IP_3$ rescues mild *par2a* mutants and restores body axis.**

**A-B:** Lateral Nomarski images of naturally fertilised 24 hpf mild *par2a* embryos injected with either KCl (A) or 20pmol $IP_3$ in KCl (B). **C:** Proportion of mild *par2a*$^{-/-}$ embryos from a single clutch failing to gastrulate or showing full or partial body axis following KCl or $IP_3$ injection. Chi-squared analysis; *** = $p<0.001$. Scale bar: B = 200µm

| Wild-type ♂ x | | Reduced Chorion Elevation | Blastodisc Absent | Defective Cell Division | Fails to Gastrulate | Total no. of embryos in clutch |
|---|---|---|---|---|---|---|
| *par2a*$^{lkc6}$ | ♀ 1 | 0% | 0% | 100% | 95% | 70 |
| | ♀ 2 | 0% | 26% | 74% | 98% | 127 |
| | ♀ 3 | 48% | 40% | 52% | 100% | 80 |
| *par2a*$^{lkc4}$ | ♀ 1 | 4% | 71% | 29% | 100% | 59 |
| | ♀ 2 | 100% | 100% | 0% | 100% | 48 |
| | ♀ 3 | 31% | 92% | 1% | 100% | 65 |
| *par2a*$^{lkc5}$; *par2b*$^{lkc7}$ | ♀ 1 | 1% | 14% | 86% | 98% | 244 |
| | ♀ 2 | 70% | 57% | 40% | 100% | 74 |
| | ♀ 3 | 0% | 67% | 32% | 100% | 100 |

**Supplementary Table 1. Penetrance of mutant phenotypes**

**Supplementary Movie 1: Inhibition of serine protease activity by Aprotinin reduces $Ca^{2+}$ wave intensity and duration during egg activation**

Fluorescent time lapse of unfertilised eggs from a *Tg(actb2:GCaMP6s)*$^{lkc2}$ female activated in E2 (left panel) or 5mg/ml Aprotinin (right panel). First 620 seconds following egg activation are shown. Scale bar = 200µm

**Supplementary Movie 2: Inhibition of serine protease activity by Benzamidine reduces $Ca^{2+}$ wave intensity and duration during egg activation**

Fluorescent time lapse of unfertilised eggs from a *Tg(actb2:GCaMP6s)*$^{lkc2}$ female activated in E2 (left panel) or 40mg/ml Benzamidine (right panel). First 620 seconds following egg activation are shown. Scale bar = 200µm

**Supplementary Movie 3: *par2a* mutant eggs show defective egg activation**

Timlapse movies of naturally fertilised (top row) and unfertilised activated (bottom row) eggs of WT (left), mild *par2a*$^{-/-}$ (middle) and severe *par2a*$^{-/-}$ (right) mutants over the first 120 minutes following fertilisation/activation. Scale bar = 200µm

**Supplementary Movie 4: *par2a* mutant embryos show failed cytoplasmic streaming**

Timlapse movies of movement of fluorescent latex beads injected into the vegetal region of WT (left) and severe *par2a*$^{-/-}$ mutants immediately following natural fertilisation. Beads were tracked (red arrowheads) over 55 minutes. Scale bar = 100µm

**Supplementary Movie 5: Mild *par2a* mutant eggs show reduced $Ca^{2+}$ wave intensity and duration during egg activation**

Fluorescent timelapse of GCaMP6s dynamics during WT (left) and mild *par2a* mutant (right) egg activation as visualised by the *Tg(actb2:GCaMP6s)*$^{lkc2}$ transgenic line. Eggs were activated in E2 and imaged for the first 695 seconds. Scale bar = 200µm

**Supplementary Movie 6: Severe *par2a* mutant eggs show strongly reduced $Ca^{2+}$ wave intensity and duration during egg activation**

Fluorescent time lapse of GCaMP6s dynamics during WT (left) and severe *par2a* mutant (right) egg activation as visualised by the *Tg(actb2:GCaMP6s)*$^{lkc2}$ transgenic line. Eggs were activated in E2 and imaged for the first 745 seconds. Scale bar = 200µm

**Supplementary Movie 7: Mild *par2a* mutant eggs show reduced $Ca^{2+}$ wave intensity and propagation at the blastomere cleavage furrow**

Fluorescent time lapse of GCaMP6s dynamics during WT (left) and mild *par2a* mutant (right) blastomere cleavage as visualised by the *Tg(actb2:GCaMP6s)*$^{lkc2}$ transgenic line. Eggs were naturally fertilised and imaged for 60 minutes. Scale bar = 200µm

**Supplementary Movie 8: $IP_3$ can partially restore $Ca^{++}$ wave intensity and duration during egg activation in severe *par2a* mutant eggs**

Fluorescent time lapse of GCaMP6s dynamics during egg activation of severe *par2a* mutants as visualised by the *Tg(actb2:GCaMP6s)*$^{lkc2}$ transgenic line. Eggs were injected with KCl alone (left) or $IP_3$ in KCl and imaged for the first 735 seconds following natural fertilisation. Scale bar = 200µm
